# Testing the efficacy of hyperspectral (AVIRIS-NG), multispectral (Sentinel-2) and radar (Sentinel-1) remote sensing images to detect native and invasive non-native trees

**DOI:** 10.1101/2021.01.01.425059

**Authors:** M Arasumani, Aditya Singh, Milind Bunyan, V.V. Robin

## Abstract

Invasive alien species (IAS) threaten tropical grasslands and native biodiversity and impact ecosystem service delivery, ecosystem function, and associated human livelihoods. Tropical grasslands have been dramatically and disproportionately lost to invasion by trees. The invasion continues to move rapidly into the remaining fragmented grasslands impacting various native grassland-dependent species and water streamflow in tropical montane habitats. The Shola Sky Islands of the Western Ghats host a mosaic of native grasslands and forests; of which the grasslands have been lost to exotic tree invasion (Acacias, Eucalyptus and Pines) since the 1950s. The invasion intensities, however, differ between these species wherein *Acacia mearnsii* and *Pinus patula* are highly invasive in contrast to *Eucalyptus globulus*. These disparities necessitate distinguishing these species for effective grassland restoration. Further, these invasive alien trees are highly intermixed with native species, thus requiring high discrimination abilities to native species apart from the non-native species.

Here we assess the accuracy of various satellite and airborne remote sensing sensors and machine learning classification algorithms to identify the spatial extent of native habitats and invasive trees. Specifically, we test Sentinel-1 SAR and Sentinel-2 multispectral data and assess high spatial and spectral resolution AVIRIS-NG imagery identifying invasive species across this landscape. Sensor combinations thus include hyperspectral, multispectral and radar data and present tradeoffs in associated costs and ease of procurement. Classification methods tested include Support Vector Machine (SVM), Classification and Regression Trees (CART) and Random Forest (RF) algorithms implemented on the Google Earth Engine platform. Results indicate that AVIRIS-NG data in combination with SVM recover the highest classification skill (Overall −98%, Kappa-0.98); while CART and RF yielded < 90% accuracy. Fused Sentinel-1 and Sentinel-2 produce 91% accuracy, while Sentinel-2 alone yielded 91% accuracy with RF and SVM classification; but only with higher coverage of ground control points. AVIRIS-NG imagery was able to accurately (97%) demarcate the Acacia invasion front while Sentinel-1 and Sentinel-2 data failed. Our results suggest that Sentinel-2 images could be useful for detecting the native and non-native forests with more ground truth points, but hyperspectral data (AVIRIS-NG) permits distinguishing, native and non-native tree species and recent invasions with high precision using limited ground truth points. We suspect that large areas will have to be mapped and assessed in the coming years by conservation managers, NGOs to plan restoration, or to assess the success of restoration activities, and several data procurement and analysis steps may have to be simplified.

## 1. Introduction

Invasive alien tree species threaten ecosystem integrity by modifying the structure and function of ecosystems and have negative impacts on ecosystem services delivery and native biodiversity (Le Maitre et al. 2011; Mooney 2005). In particular, three genera of invasive alien trees - acacias, pines and eucalyptus species in South Africa (Gaertner et al. 2017; McConnachie et al. 2015), Brazil (de Abreu, Durigan 2011), (Argentina Zalba et al. 2008), Uruguay (Six et al. 2013), (Kenya Pellikka et al. 2009), New Zealand (Ledgard 2001), Hawaiian Islands (Daehler 2005) and India (Arasumani et al. 2019) are often listed as the worst offenders. These trees are native to Australia and were introduced to several tropical and subtropical countries in the nineteenth century (Richardson 1998). These trees were largely established on grasslands and shrublands as these landscapes were historically classified as wastelands (Joshi et al. 2018; Rundel et al. 2014). All of these species are fast-growing, have high water use and show potential for encroachment into native grasslands and scrublands. Acacias, in particular form dense stands, maintain a high leaf area all through the year and have high evapotranspiration which impacts water yields of infested catchments (Dye, Jarmain 2004). These species have also had negative impacts on grasslands birds of South Africa (Allan et al. 1997; Armstrong, Van Hensbergen 1995), on small mammals (Armstrong, Van Hensbergen 1995), invertebrates (Donnelly 1985) and plants (Richardson, Van Wilgen 1986).

In the Western Ghats, tropical montane grasslands, also known as Shola grasslands have been lost to exotic trees invasion at a rapid pace (Arasumani et al. 2018; Arasumani et al. 2019). This widespread invasion has impacted various faunal communities, including threatened species like the Nilgiri Pipit (Lele et al. 2020) and Nilgiri Tahr (Alempath 2008) in the Western Ghats. These trees were primarily established outside wildlife preserves and protected areas since the 1950s. However, today, the invasion of exotic trees is rapidly encroaching critical grassland landscapes, including protected areas (Arasumani et al. 2019; Joshi et al. 2018). Our prior research using Landsat data (Arasumani et al. 2019) indicates that 23% (340 Km2) of montane grasslands have been exotic tree stands within the past five decades. This data, however, had included all exotic trees as a single class due to the limitations with the spectral and spatial resolution of LANDSAT imagery. Single-class invasive species maps are however limiting as invasion by exotic species occurs at different speeds, with some species (mainly acacias and pines) rapidly invading into remnant fragmented montane grasslands in the Western Ghats (Arasumani et al. 2019). These invasive trees - *Acacia mearnsii* and *Pinus patula* have been listed among the most invasive species in the World (IUCN-GISD), but they co-occur with less invasive species as well as native tropical forests (Joshi et al. 2018) making the detection and estimation of the spread of the invasive trees challenging. One of the major challenges for conservation managers is to detect the invasion front, which typically consists of sparsely dispersed seedlings and saplings that are difficult to detect using remote sensing imageries (but see Arasumani et al. 2020). Detecting this invasion front is, however, critical in managing the invasion front. In addition, to identify the best combination of algorithms and remote sensing platforms that can be used and accessed by conservation agencies in tropical areas across the globe.

### 1.1. Choice of imageries

Field-based, landscape-scale, invasive species mapping techniques are known to be challenging in tropical forests. While many researchers have attempted to identify broad forest type categories using space-borne and airborne images (Arasumani et al. 2019; Erinjery et al. 2018; Foody, Hill 1996; Shimizu et al. 2019), discriminating between native and non-native species is challenging with medium resolution satellite data due to extensive intermixing and canopy heterogeneity requires high spatial and spectral resolution imagery. Researchers have also employed microwave data (Chen et al. 2018; Laurin et al. 2013; Wheeler et al. 2017), recently launched Sentinel-2 multispectral images (Laurin et al. 2013; Wheeler et al. 2017), and combinations of Sentinel-1 SAR (Synthetic Aperture Radar) images with Sentinel-2 multispectral images to improve classification accuracy (Erinjery et al. 2018; Kattenborn et al. 2019; Zhang et al. 2019). A few studies have also suggested that hyperspectral images could produce high accuracy for mapping tree species compared to the multispectral datasets (George et al. 2014; Thenkabail et al. 2004).

### 1.2. Choice of classification algorithms

Several classification algorithms have been used for classifying hyperspectral, multispectral and SAR data: of these, Support Vector Machine (SVM) (George et al. 2014), Random Forest (RF) and Classification and Regression Tree (CART) have been the most widely used especially on the publicly-available Google Earth Engine Platform (Gorelick et al. 2017). Although some studies pick the best classification algorithm for classifying hyperspectral data; all three algorithms are used for landcover mapping using both multispectral and SAR images (Lu et al. 2018) with perhaps differences in their training data requirements. The SVM is an iterative, non-parametric machine learning algorithm widely used for classifying the hyperspectral images (Mountrakis et al. 2011). The SVM does not depend on the statistical distribution of the data but relies on training data adjacent to the class boundary to deliver high accuracy even with limited training data for classification (Melgani, Bruzzone 2004). The random forest (RF) is a non-parametric classifier that operates by generating a number of classification trees and selecting the mode of the predictions (Rodriguez-Galiano et al. 2012). CART models use recursive binary splits on predictor data in a decision tree framework to produce classifications at the end nodes of the trees. By nature of the classification process, CART models are considered somewhat easier to interpret compared to RF-based models (Lawrence, Wright 2001). Notably, RF and CART approaches are widely used to classify the remote sensing images but have higher training data requirements relative to SVM models (Delalay et al. 2019; Shaharum et al. 2020).

### 1.3. Objectives

Overall, this study aims to identify appropriate data sources and algorithms to identify exotic tree species on the Sky Islands of the Western Ghats. Specifically, we test the ability of a) AVIRIS-NG (hyperspectral), Sentinel-2B (multispectral) and Sentinel-1B (microwave) data with b) SVM, Random Forest, and CART classification algorithms to discriminate invasive woody species (Acacia, Pine and Eucalyptus) from native tropical trees in the Shola Sky Islands. The overall goal is to test the efficacy of these imageries in detecting the invasion front - the ecotone where conservation efforts can be targeted.

## 2. Methods

### 2.1. Study area

For the purposes of this study, we selected an area admeasuring approximately 12 sq. km. in the Nilgiris mountains (Fig. 1) that has a gradient of woody invasive species infestation across this landscape. The study area contains natural habitats such as montane grasslands, montane forests and water bodies, and non-native woody trees - *Acacia spp, Eucalyptus spp*, and *Pinus spp* (Fig. 2).

**Fig. 1.**
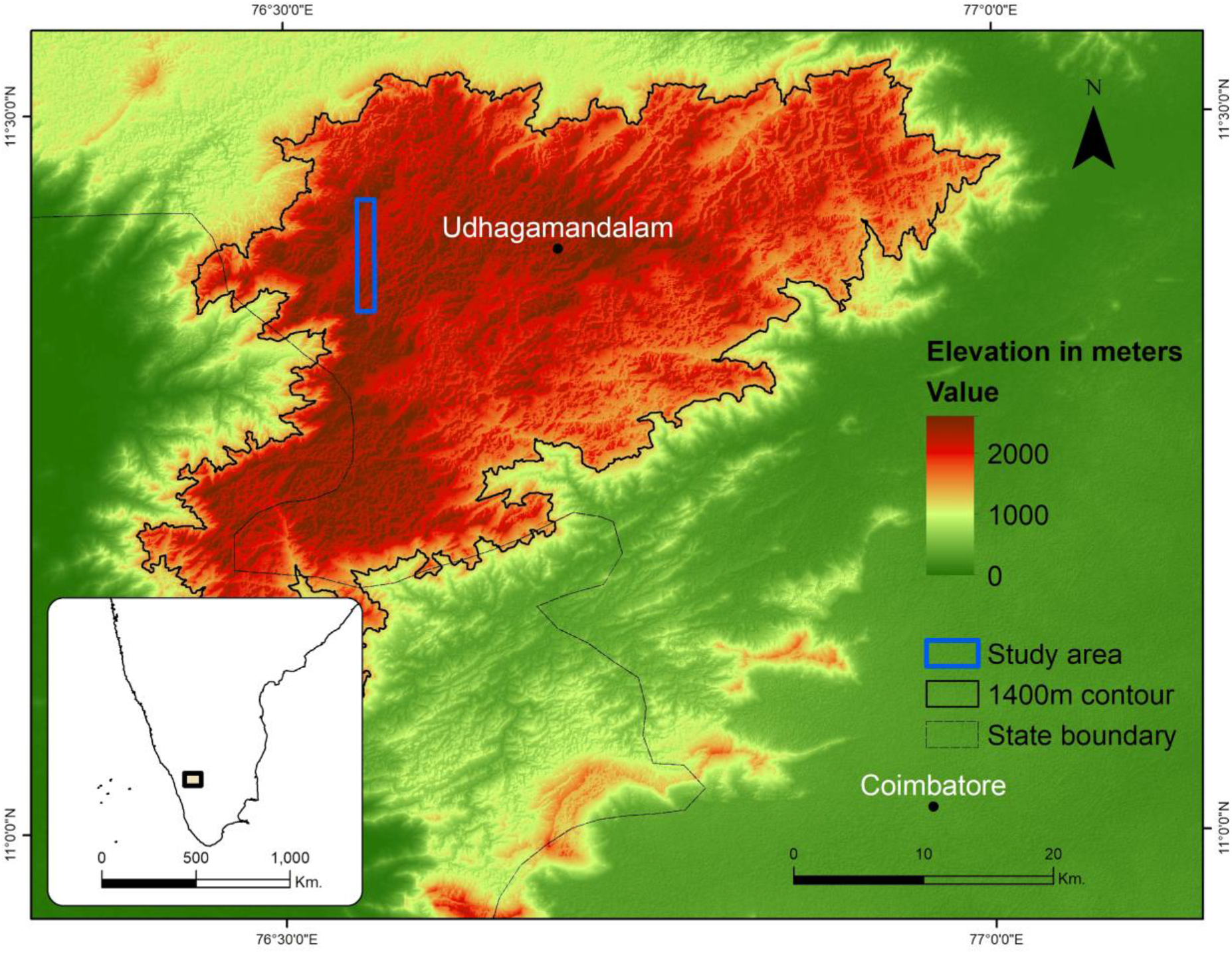
Study area - Nilgiri Hills.

**Fig. 2.**
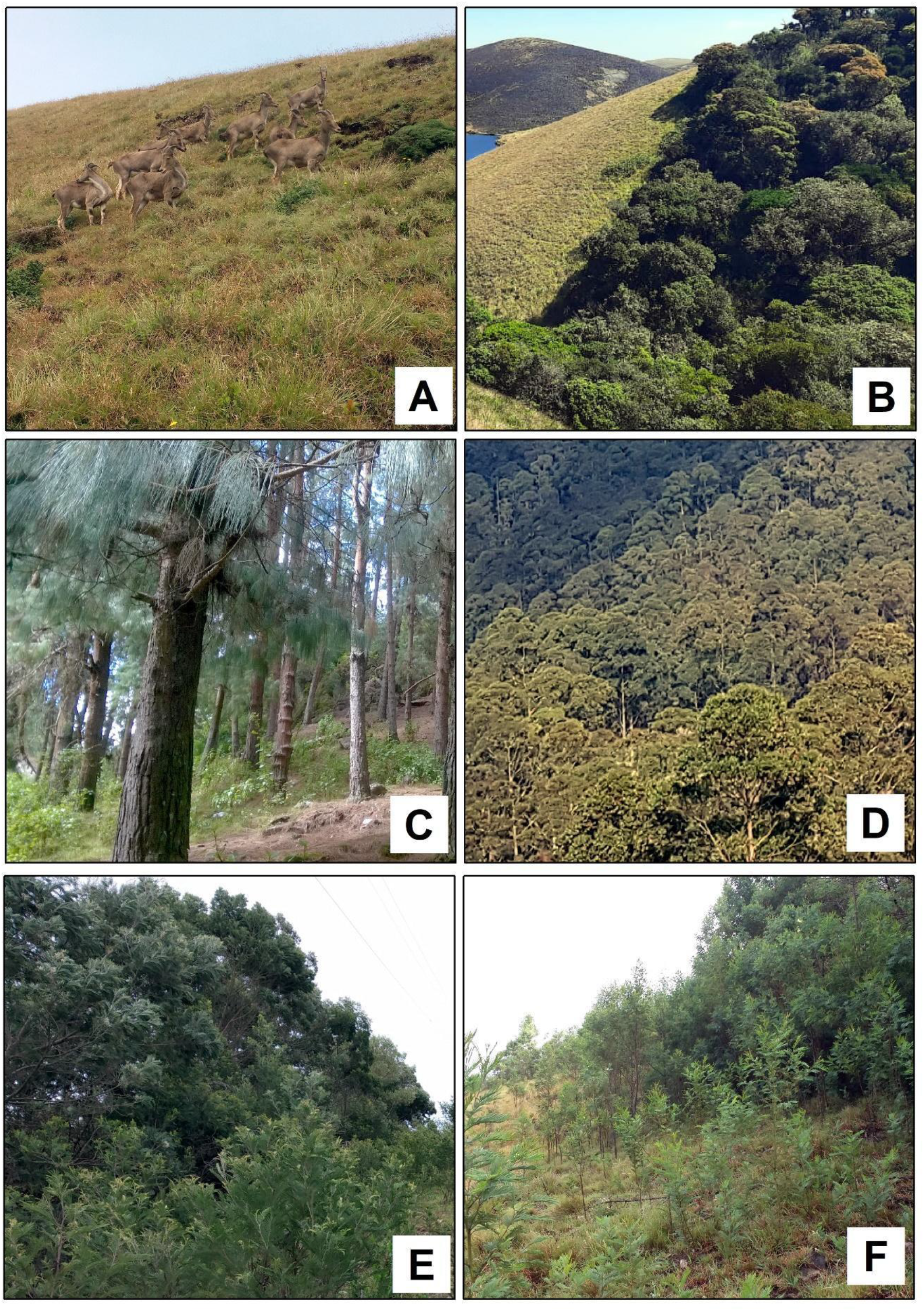
Field photographs; A – Montane Grasslands, B – Montane forests, C - Pine, D – Eucalyptus, E - Acacia, and F – Acacia invasion front.

### 2.2. Field data

In our study region, native forests are generally intermixed with non-native trees (*Acacia spp*., *Pinus spp*., and *Eucalyptus spp*.). However, only acacias were observed to be present at the actual grassland-forest invasion front (i.e. pines and eucalypti were generally established in distinct stands). We collected the unique land cover GPS points where target species land covered in excess of a 30m*30m footprint to obtain ‘pure’ endmember spectral signatures. We used 1049 ground-truth points of montane grasslands, forest, acacias, eucalyptus, pines and water bodies for image classification. For all locations, we ensured that the positional dilution of precision (PDOP) was lower than the AVIRIS-NG, Sentinel-1, and Sentinel-2-spatial resolutions (4m and 10m). To detect the *Acacia spp*. invasion front, we collected 73 additional ground truth locations concurrent with satellite imagery. We assessed the efficiency of acacia invasion front mapping with classifications produced by the three different image datasets and algorithms on Google Earth Engine Platform as described below.

### 2.3. Image Data

We obtained an AVIRIS-NG (Airborne Visible InfraRed Imaging Spectrometer - Next Generation) apparent at-surface reflectance products from the Jet Propulsion Laboratory (JPL), National Aeronautics and Space Administration data portal (NASA - https://avirisng.jpl.nasa.gov/dataportal/). AVIRIS-NG data have high spatial (4 m) and spectral (5 nm) spectral resolution with 425 spectral bands spanning 380 nm – 2510 nm. We excluded noisy bands and water vapour absorption bands (bands 1-10, 195-207, 287-316, 325-329) from the AVIRIS-NG dataset.

The Sentinel-2 Level-2 ground reflectance product was not available for our study area on the GEE platform for 2018. We, therefore, substituted this with the Sentinel-2 Level-1 product from the USGS (United States Geological Survey) Earth Explorer portal (https://earthexplorer.usgs.gov/), and we performed atmospheric corrections using Sen2Cor v2.8 to convert at-sensor radiance imagery to apparent at-surface reflectance. We standardized all bands to a 10m spatial resolution for all subsequent analyses.

We obtained the Sentinel-1 SAR Ground Range Detected products directly from the GEE platform. This data was available as calibrated and ortho-corrected and was pre-processed using the Sentinel-1 toolbox for thermal noise removal, radiometric calibration, and terrain correction (using the SRTM 30 m spatial resolution digital elevation model). The final terrain-corrected data were log-transformed to decibels. We used Sentinel-1 VH (vertical transmit and horizontal receive) and VV (vertical transmit and vertical receive) polarization for image classification. We fused the Sentinel-1 (VV, VH bands) and Sentinel-2 (2, 8) bands for all subsequent analyses.

### 2.4. Classification algorithms

All image classifications were conducted using the Google Earth Engine (GEE) platform. We chose GEE to enable the creation of a workflow that can be utilized by conservation managers in other regions to track and manage invasions.

We tested a combination of Random Forest (RF; Breiman 2001), Classification and Regression Tree (CART; Breiman et al. 1984), and Support Vector Machine using radial basis functions (SVM; Burges 1998) to assess the skill of classification. For the SVM, parameter values (gamma and cost) were determined by using an iterative grid search.

We assessed the accuracy of the classifications using 300 ground truth points held out from all preceding analyses and estimated accuracy metrics (overall accuracy, user accuracy, producer accuracy and, Kappa coefficient) from the confusion matrix (Congalton 1991).

## 3. Results

### 3.1. Comparison of different classification

We observed the highest overall accuracy (OA; 98.7%) and kappa coefficient (Kappa; 0.984) in AVIRIS-NG dataset with SVM classification (Table 1, Fig. 3), followed by S1+S2 datasets with CART (OA − 91.3%, Kappa − 0.896), Sentinel-2 with SVM (OA − 91.0%, Kappa − 0.892) and Sentinel-2 with RF (OA − 91.0%, Kappa − 0.892)

**Table 1.**
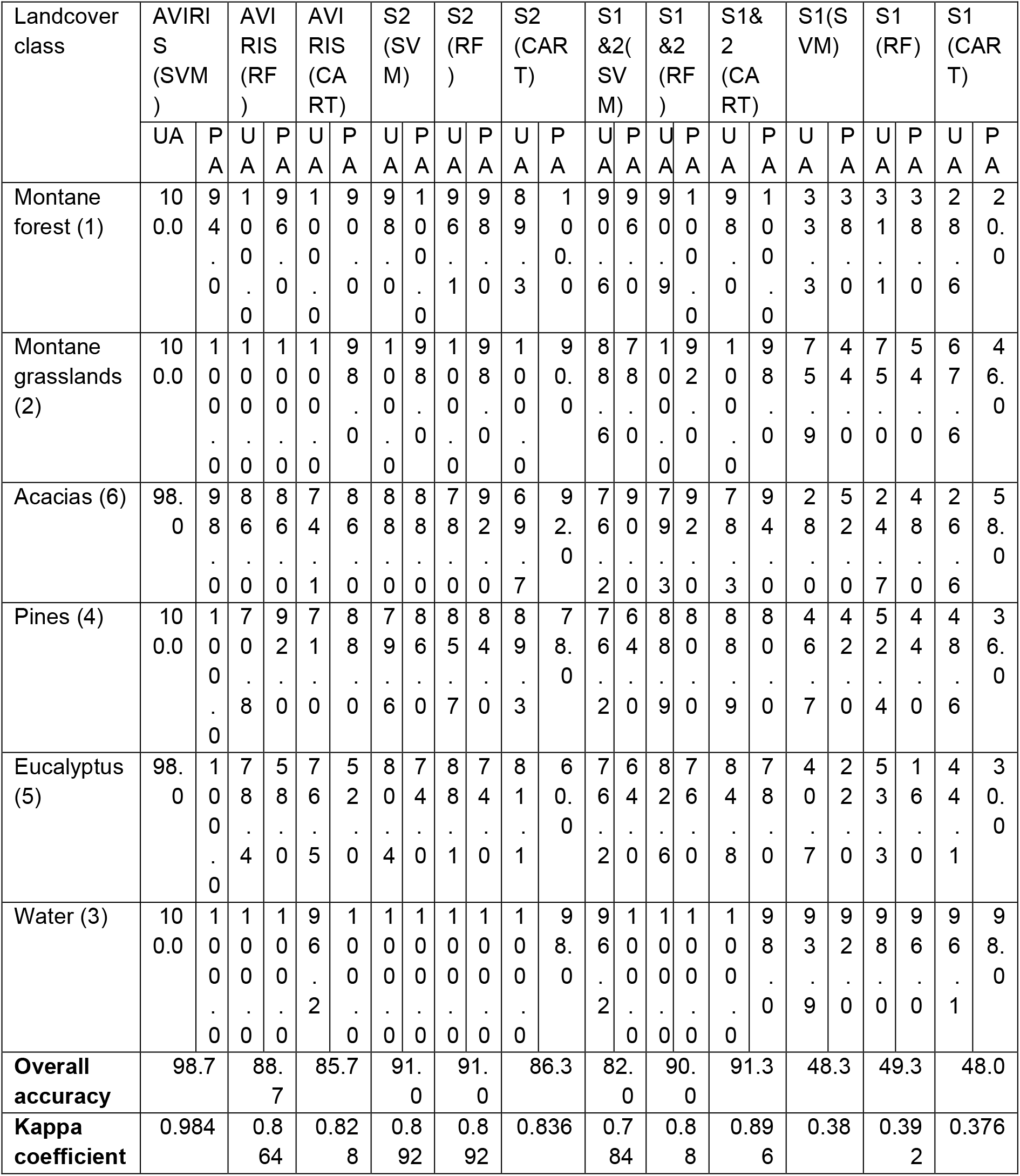
Image classification results. S1 - Sentinel-1; S2 - Sentinel-2; UA-User Accuracy; PA-Producer Accuracy.

**Fig. 3.**
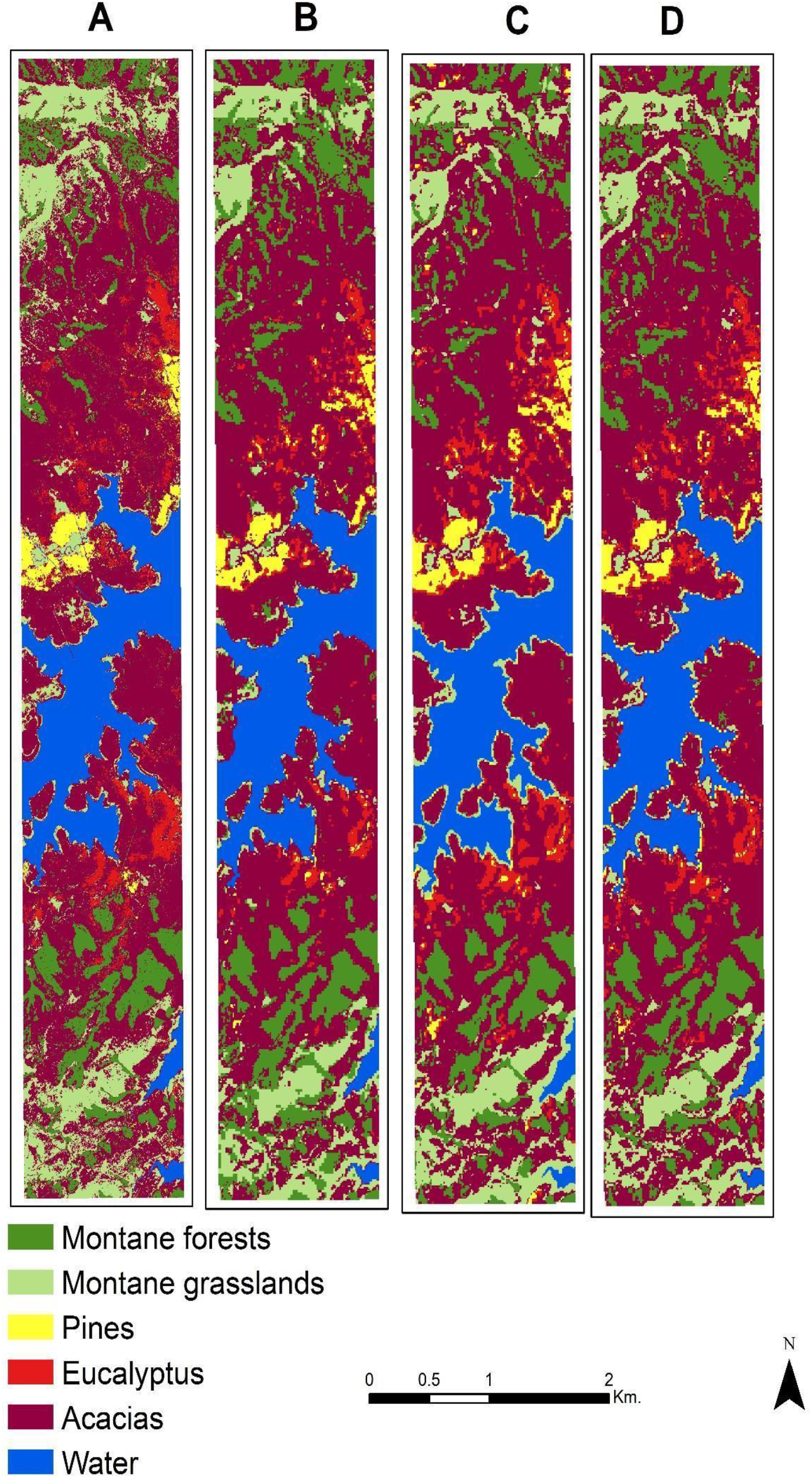
**The best-classified maps of native and non-native habitats. A – AVIRIS-NG with SVM, B – S1+S2 datasets with RF, C - Sentinel-2 with SVM, and D – Sentinel-2 with RF**.

The highest overall accuracy in the AVIRIS-NG dataset was recorded with SVM classification (OA − 98.7, Kappa − 0.984), followed by Random Forest classification (OA − 88.7 Kappa − 0.864) and CART classification (OA − 85.7 Kappa − 0.828). (Table 1, Fig. 4).

**Fig. 4.**
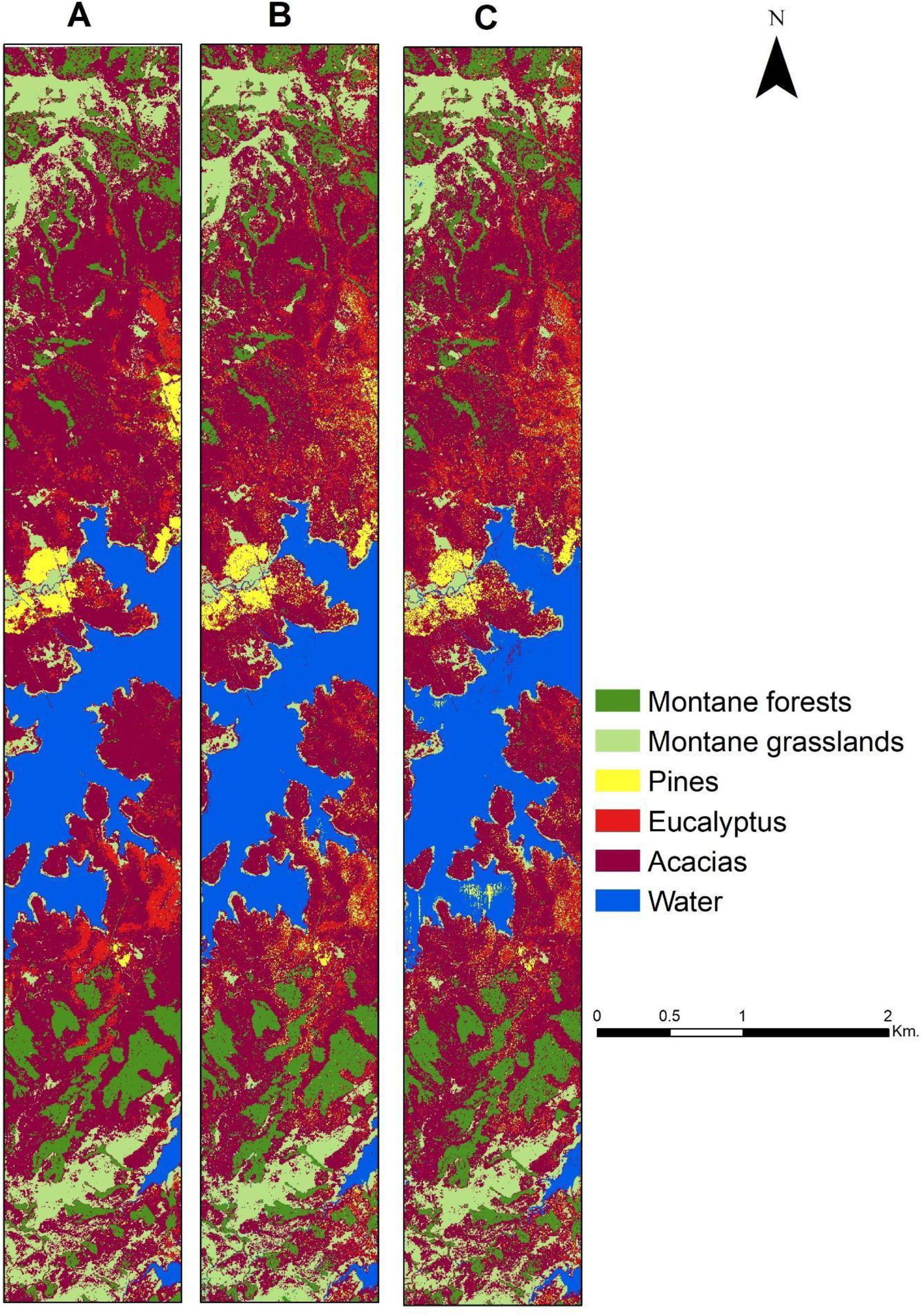
The AVRIS-NG classified maps of native and non-native habitats. A – Classified with SVM, B – RF, C – CART.

The highest overall accuracy for Sentinel-2 data was also obtained with SVM classification (OA − 91.0, Kappa − 0.892) and RF classification (OA − 91.0, Kappa − 0.892), followed by CART classification (OA − 86.3, Kappa − 0.836) with S2 bands of 2,3,4,5,6,7 and 8 (Table 1, Fig. 5).

**Fig. 5.**
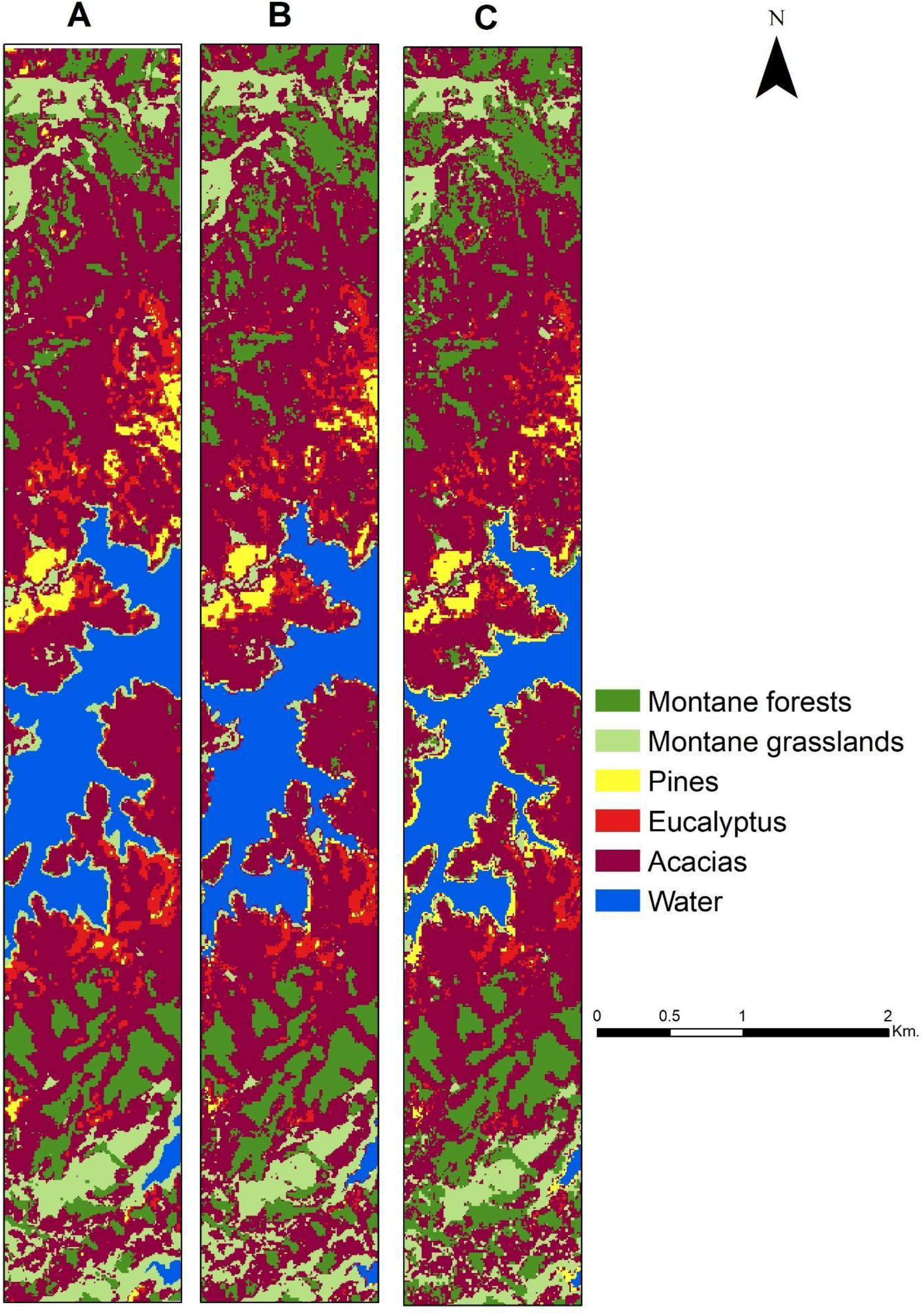
The Sentinel-2 classified maps of native and non-native habitats. A – Classified with SVM, B – RF, and C – CART.

The S1+S2 datasets provided the highest overall accuracy with CART classification (OA − 91.3, Kappa − 0.896) and RF classification (OA − 90.0, Kappa − 0.88), followed by SVM classification (OA − 82.0, Kappa − 0.784) with Sentinel-2 bands 2,3,4,5,6,7,8 and Sentinel-1 VV and VH polarisations (Table 1, Fig. 6).

**Fig. 6.**
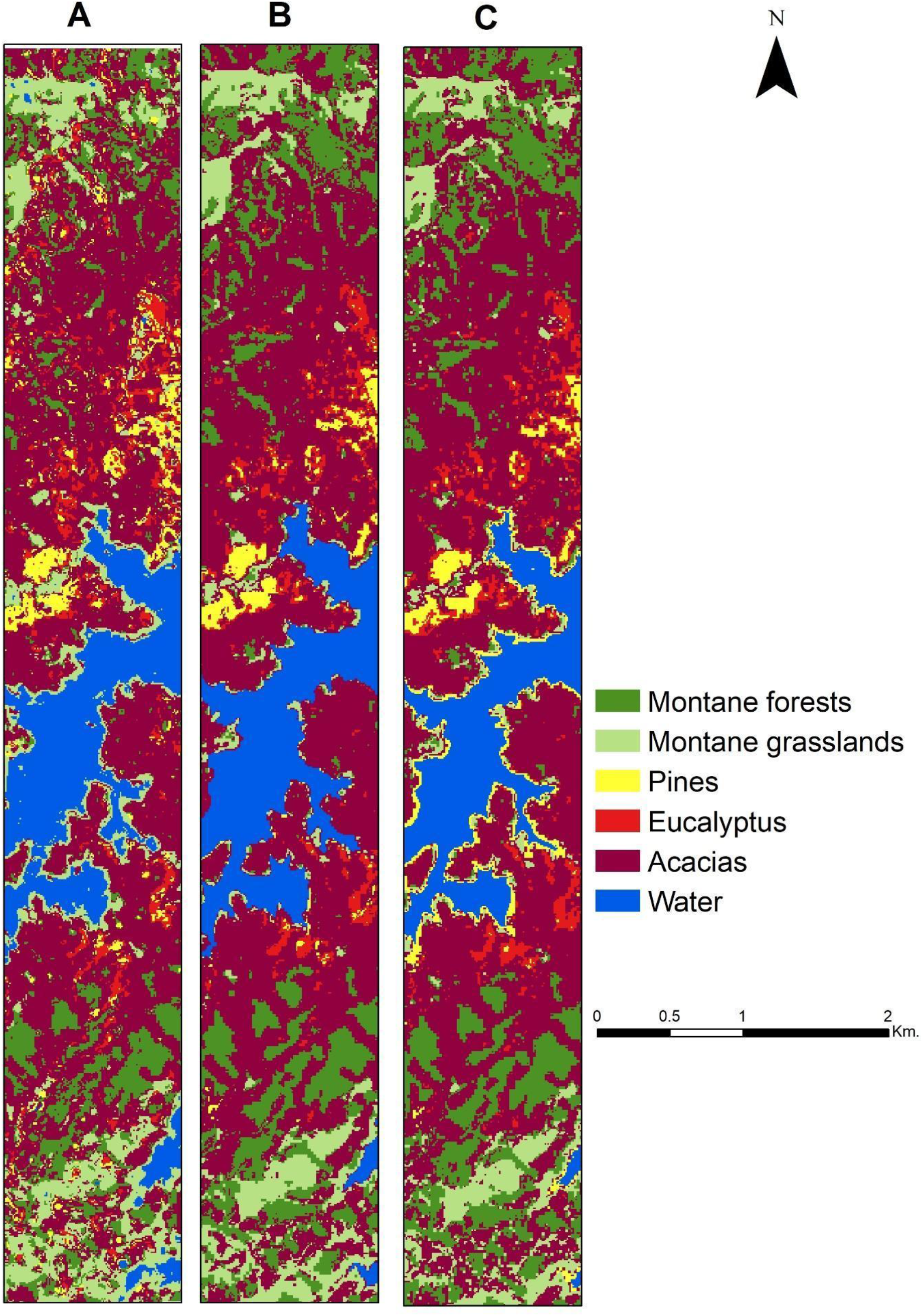
The Sentinel-1 and Sentinel-2 classified maps of native and non-native habitats. A – Classified with SVM, B – RF, and C – CART.

The Sentinel-1 dataset resulted in low accuracy of all classifiers compared to the other datasets with the highest overall accuracy with RF classification (OA − 49.3, Kappa-0.392), and CART classification (OA − 48.0, Kappa-0.376), followed by SVM classification (OA − 48.3, Kappa-0.38) with VV and VH polarisations (Table 1, Fig. 7).

**Fig. 7.**
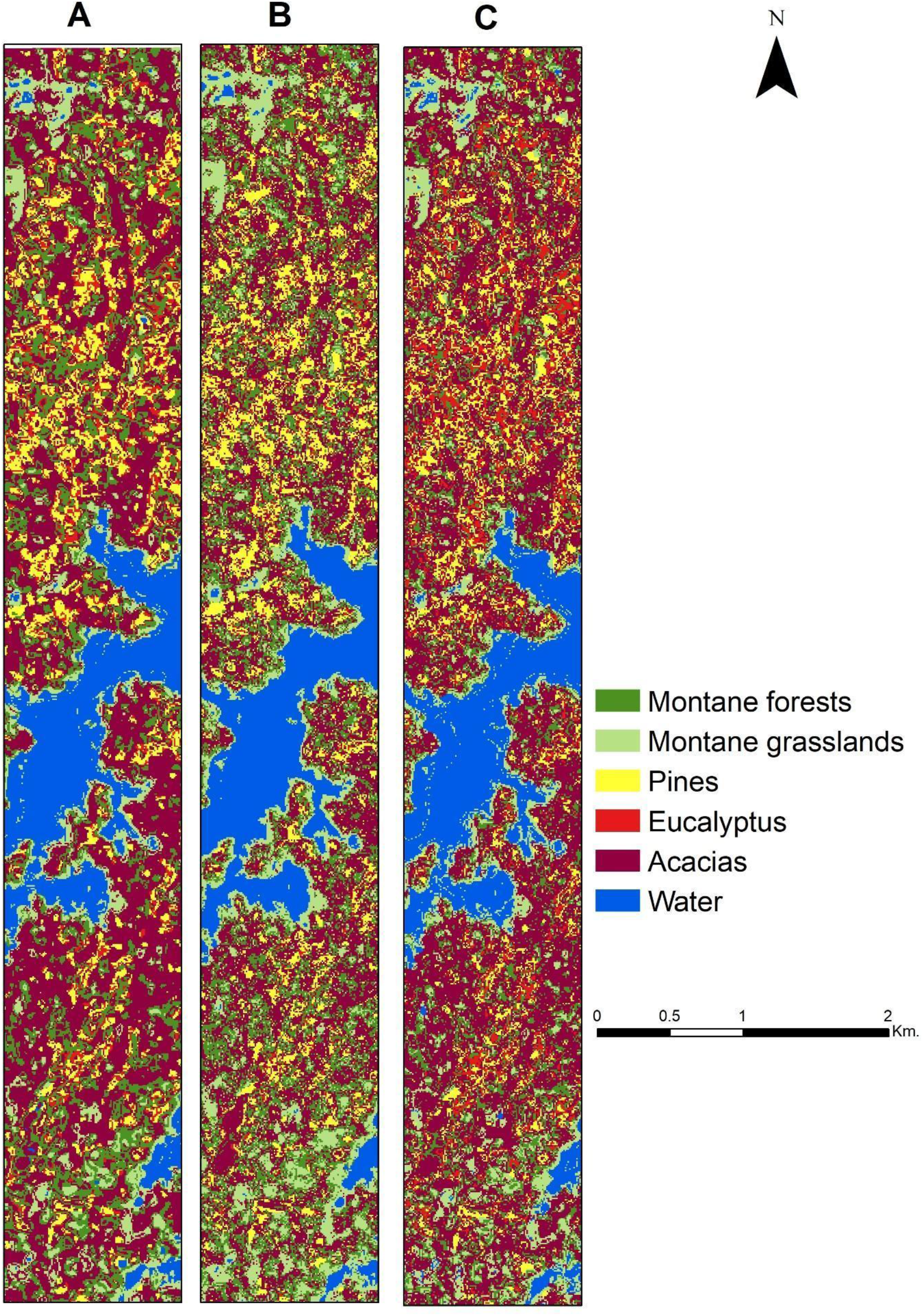
The Sentinel-1 classified maps of native and non-native habitats. A – Classified with SVM, B – RF, and C – CART.

### 3.2. Best producer & user accuracy

The highest producer accuracy and user accuracy were obtained from AVIRIS-NG dataset with SVM of all classes. The Sentinel-2 with SVM and S1+S2 datasets with CART classifications produced high producer & user accuracy in detecting in water, forests and grasslands; however, it produced marginally lower efficiency in identifying acacias, eucalyptus and pine. However, Sentinel-1 performed poorly, and the user and producer accuracy of all the classes was low (<50%) except for water (Table 1).

### 3.3. Number of training samples vs classification accuracy

The AVIRIS-NG dataset with SVM classification produced 94% of overall accuracy and 0.93 kappa coefficient with limited data (25% of training samples) however RF and CART classifications yielded less than 81% of overall accuracy (Table 2). The Sentinel-2 with RF classification produced overall accuracy 89% and kappa coefficient 0.85 with 25% of training samples. We observed that AVIRIS-NG dataset with SVM classification produced very high accuracy (97%) with the 50% per cent of training samples; however, Sentinel-2 and S1+S2 datasets yielded less than 90% overall accuracy. We noted that the classification accuracy for S1+S2 datasets did not vary much with partial training datasets of 75% when compared to the complete training data set (Table 2).

**Table 2.** Accuracy assessment with 25%, 50%, 75% and 100% of training samples. UA-User Accuracy; PA-Producer Accuracy.

|  | OA<br>(25%) | Kappa<br>(25%) | OA<br>(50%) | Kappa<br>(50%) | OA<br>(75%) | Kappa<br>(75%) | OA<br>(100%) | Kappa<br>(100%) |
| --- | --- | --- | --- | --- | --- | --- | --- | --- |
| AVIRIS-NG<br>(SVM) | 94.00 | 0.93 | 97.00 | 0.96 | 97.67 | 0.97 | 98.67 | 0.98 |
| AVIRIS-NG<br>(RF) | 81.33 | 0.78 | 85.67 | 0.83 | 86.33 | 0.84 | 88.67 | 0.86 |
| AVIRIS-NG<br>(CART) | 73.00 | 0.68 | 84.00 | 0.81 | 85.33 | 0.82 | 85.67 | 0.83 |
| Sentinel-2<br>(SVM) | 80.33 | 0.76 | 89.67 | 0.88 | 91.33 | 0.90 | 91.00 | 0.89 |
| Sentinel-2 (RF) | 89.00 | 0.87 | 90.67 | 0.89 | 91.00 | 0.89 | 91.00 | 0.89 |
| Sentinel-2<br>(CART) | 87.67 | 0.85 | 89.00 | 0.87 | 89.67 | 0.88 | 86.33 | 0.84 |
| Sentinel-1&2<br>(SVM) | 65.67 | 0.59 | 72.67 | 0.67 | 80.00 | 0.76 | 82.00 | 0.78 |
| Sentinel-1&2<br>(RF) | 89.33 | 0.87 | 89.00 | 0.87 | 90.67 | 0.89 | 90.00 | 0.88 |
| Sentinel-1&2<br>(CART) | 87.67 | 0.85 | 90.33 | 0.88 | 88.33 | 0.86 | 91.33 | 0.90 |

### 3.4. Detecting the acacia invasion front

We observed high accuracy in detecting Acacia invasion front with AVIRIS-NG dataset with SVM classification (97%). The RF and CART classification, however, produced lower accuracy (<40%). Moderate accuracy was observed in the Sentinel-2 with RF classification (60%) followed by SVM classification (58%) and CART classification (38%). The S1+S2 datasets and Sentinel-1 alone produced less than 40% accuracy across all classifiers.

## 4. Discussion

Systematic monitoring and mapping of invasive alien species are essential for conservation and restoration of tropical grasslands. Spatially-explicit information on native habitats and non-native species is critical for sustainable forest management and forecasting landscape changes into the future.

### 4.1. Image data sources

In this study, we found that the hyperspectral dataset (AVIRIS-NG in this study) was the ideal platform for discriminating between native and non-native invasive tree species with high precision. The AVIRIS-NG dataset accurately delineated the edges of non-native trees and native habitats, likely due to its high spatial and spectral resolution. The S1+S2 datasets comprising multispectral and radar data proved to be a reasonable alternative but were only marginally better than using Sentinel-2 images alone. Finally, Sentinel-1 data did not produce sufficient accuracy for classifying and differentiating the invasive species from the native species in the tropical montane habitats. Similar results have been reported in classifying forest types with Sentinel-1 data in these tropical regions (Erinjery et al. 2018).

Our study supports results obtained by others using hyperspectral remote sensing data for tropical tree species mapping efforts (Hyperion data − 30 m spatial resolution and 10 nm spectral resolution; George et al. 2014; Thenkabail et al. 2004). While Sentinel-1 images have shown to be useful for detecting and classifying water bodies in numerous studies (e.g. Bioresita et al. 2018; Hu et al. 2020), we were unable to map invasive woody species accurately. This may be due to similarities in the backscattering SAR signatures of native and non-native species. Sentinel-1 also has a shorter wavelength (C-band) that may not be able to differentiate the forest types based on the height and volume information where longer wavelengths (L-band) may be of advantage (Mitchell et al. 2014).

Although our study reports a relatively high accuracy of the combined Sentinel-1 and Sentinel-2 imageries, the difference with the use of only Sentinel-2 was marginal. This, however, is different from other studies that have reported relatively higher accuracy in the classification from the combined Sentinel-1 and Sentinel-2 imageries for mapping different forest types (Erinjery et al. 2018; Kattenborn et al. 2019), agricultural applications (Veloso et al. 2017), mapping wetlands (Slagter et al. 2020), extracting urban extents (Iannelli, Gamba 2019), and delineating water bodies (Ahmad et al. 2019).

### 4.2. Classification algorithms

SVM produced the highest accuracy for classifying the hyperspectral data contrast to RF and CART. This is in agreement with several other studies that have also observed SVM to be highly suitable for classifying hyperspectral data (Burai et al. 2015; George et al. 2014; Lim et al. 2019; Raczko, Zagajewski 2017). Both RF & SVM algorithms produced good classification accuracy with Sentinel-2 images compared to CART, as also observed by Lu et al. (2018). In our tests, we observe that CART shows reasonable overall accuracy, but some grassland and acacia invasions around the water bodies tended to be misclassified with pine trees; this was also the issue with a S1+S2 classification. Although SVM performed well with Sentinel-2 data (91%), it produced low accuracy for the classification with the S1+S2 datasets (82%). This is perhaps because SVM does not do well with noisy data typical to Sentinel-1 data, and when target land cover class may have similar backscattered textures.

While Sentinel-2 did fairly well in detecting invasives and natural habitats, we found that the combination of AVIRIS-NG data with an SVM classification model was the only sensor to detect acacia invasion-front with high accuracy. While we are not aware of other studies focusing on detecting invasion fronts, AVIRIS-NG data have been successful for mapping mangroves to the species-level (Chaube et al. 2019) and for crop type identifications (Salas et al. 2020).

In this research, we utilized medium (S1 & S2 − 10m) and high spatial resolution (AVIRIS-NG − 4m) images for IAS mapping. We were therefore successful in mapping the species at the pixel scale. We suspect that sub-pixel classification methods may be more suitable for data with spatial resolutions images over 30m.

### 4.3. Influence of the number of ground truth points

We observed that a limited number of ground truth points (∼25%) were sufficient for obtaining high accuracies when using hyperspectral data (AVIRIS-NG). However, multispectral data (Sentinel-2) seemed to require a higher density of ground truth points (> 50%) to get reasonable accuracies. If the study area is large, Sentinel-2 data might need three times the number of ground truth points than that required to classify a similar hyperspectral image. Conversely, hyperspectral data (AVIRIS-NG) is not available for all places, but data can be captured using a UAV-borne hyperspectral sensor. Using an UAV, however, might require significant effort and investment for large study regions.

### 4.4. Influence of spatial and spectral resolution

We believe that the higher spatial resolution of AVIRIS-NG dataset over S1 & S2 provides a distinct advantage in distinguishing smaller native forest patches from non-native trees. We found that the S1&S2 datasets were not able to detect acacia invasion due to the lower spatial resolution compared to AVIRIS-NG dataset.

The fine spectral resolution of AVIRIS-NG (spanning 425 spectral bands) compared to Sentinel-2 (8 spectral bands) helped discriminate between individual native and invasive species. Other studies have shown similarly high species-level classification accuracies in complex landscapes (George et al. 2014; Skowronek et al. 2017). Spectral signature overlaps in the Sentinel-2 dataset resulted in lower average accuracy in detecting pines and eucalyptus similar to findings from Pu et al. (2012) and Marshall, Thenkabail (2015).

### 4.5. Inferences for conservation managers

Mapping and distinguishing native trees from non-native trees is an essential task for land managers for conserving native and endemic species, assessing fire risks, and impacts on ecosystem services. We found that airborne hyperspectral imagery provides the best solution for detecting invasive species and the invasion front. However, the synoptic availability of hyperspectral images is a major limiting factor for most locations. These data were only available only for 12 sq. km. of our study area. Further, procuring such data can be prohibitively expensive. Where these data are not available, we recommend using Sentinel-2 satellite data with SVM or RF algorithms as it offers a reasonable compromise between accurately discriminating native and non-native trees while allowing the mapping of large spatial extents. Sentinel-2 images, however, require a high-density of ground truth points, and may still be unsuitable for mapping young invasion because of the limited spatial and spectral resolution. Conservation managers may also consider using RapidEye images with an object-oriented classification approach if they need to map a mixed-species invasion front, and do not need to discriminate among the invasive species along that front (Arasumani et al. 2020). If the invasion front requires constant monitoring in a smaller study area, an Unmanned Aerial Vehicle with a hyperspectral sensor may be indispensable. Finally, we also recommend that conservation managers and restoration NGOs use the online Google Earth Engine (GEE) platform because of the high processing power required for processing high spatial and spectral resolution data. We provide the GEE code written for this study in Appendix 1. We have detailed our recommendations for conservation managers in Table 3.

**Table 3.** Recommendations for Conservation managers.

| Data | Best classification algorithms | Possible Applications | Advantages | Limitations |
| --- | --- | --- | --- | --- |
| Hyperspectral data (AVIRIS-NG or UAV with Hyperspectral sensor) | SVM | 1. Invasive species mapping.<br>2. Forest species mapping.<br>3. Invasion front detection. | 1. Require limited ground truth points for classification.<br>2. High precision outputs. | 1. Applicable for smaller areas.<br>2. Presently, airborne and satellite data are not available for all the areas.<br>3. Capturing and processing UAV data might be time-consuming.<br>4. The requirement to upload the data into Google Earth Engine. |
| Multispectral data (Sentinel-2) | SVM & RF | 1. Invasive species mapping. | 1. Data are available globally.<br>2. Data already available in Google earth engine.<br>3. Suitable for larger areas. | 1. Needs a high-density of ground truth points.<br>2. Limited accuracy compared to hyperspectral data.<br>3. Difficult to classify small patches of forests.<br>4. Invasion front mapping not possible at species level. |
| Fused dataset - Radar and multispectral (S1+S2) | RF | 1. Invasive species mapping. | 1. Data are freely available in GEE globally. | 1. Needs more ground truth points.<br>2. Limited accuracy.<br>3. Radar data contains backscatter noise or similar backscatter signature for all the species.<br>4. Invasion front mapping not possible at species-level. |
| Radar data (Sentinel-2) | RF | 1. Water bodies extraction. | 1. Data are freely available in GEE across the world.<br>2. Water bodies can be extracted during the monsoon season. | 1. Invasive species and forest mapping are not possible. |

In this research, we demonstrate the advantage of remotely-sensed hyperspectral, multispectral, and radar data for mapping, and distinguishing native and non-native invasive species using AVIRIS-NG, Sentinel-1 and Sentinel-2 datasets. Although high-resolution, hyperspectral AVIRIS-NG dataset proved superior, we were severely constrained by its spatial availability. The multispectral Sentinel-2 data, on the other hand, were useful in identifying native and non-native trees across a large landscape. Combined with the processing power of the GEE platform, this study demonstrates the opportunity for species-specific discrimination of invaded landscapes, that can be replicated across the globe.

## Acknowledgements

This research was supported by an IISER-Tirupati postdoctoral research fellowship to Arasumani M and a grant from the Ministry of Environment, Forest and Climate Change, Government of India (grant no. F. No. 19-22/2018/RE) to V.V. Robin. We thank Tamil Nadu forest department and APCCF & Field Director, Mudumalai Tiger Reserve for logistics and research permits. We thank Vasanth Bosco for extensive discussions and feedback. We thank Ecology and Evolution lab members at IISER-Tirupati for comments and feedback on the manuscript.

## References

Ahmad SK, Hossain F, Eldardiry H, et al. (2019) A Fusion Approach for Water Area Classification Using Visible, Near Infrared and Synthetic Aperture Radar for South Asian Conditions. IEEE Transactions on Geoscience and Remote Sensing

Alempath MR, C. 2008 (2008) Nilgiritragus hylocrius. In.

Allan DG, Harrison JA, Navarro R, et al. (1997) The impact of commercial afforestation on bird populations in Mpumalanga Province, South Africa—insights from bird-atlas data. Biological Conservation 79:173–185

Arasumani M, Bunyan M, Robin VV (2020) Opportunities and challenges in using remote sensing for invasive tree species management, and in the identification of restoration sites in tropical montane grasslands. Journal of Environmental Management:111759

Arasumani M, Khan D, Das A, et al. (2018) Not seeing the grass for the trees: Timber plantations and agriculture shrink tropical montane grassland by two-thirds over four decades in the Palani Hills, a Western Ghats Sky Island. PLOS ONE 13:e0190003

Arasumani M, Khan D, Vishnudas C, et al. (2019) Invasion compounds an ecosystem-wide loss to afforestation in the tropical grasslands of the Shola Sky Islands. Biological conservation 230:141–150

Armstrong A, Van Hensbergen H (1995) Effects of afforestation and clearfelling on birds and small mammals at Grootvadersbosch, South Africa. South African Forestry Journal 174:17–21

Bioresita F, Puissant A, Stumpf A, et al. (2018) A method for automatic and rapid mapping of water surfaces from sentinel-1 imagery. Remote Sensing 10:217

Breiman L (2001) Random forests. Machine learning 45:5–32

Breiman L, Friedman J, Stone CJ, et al. (1984) Classification and regression trees. CRC press

Burai P, Deák B, Valkó O, et al. (2015) Classification of herbaceous vegetation using airborne hyperspectral imagery. Remote Sensing 7:2046-2066

Burges CJ (1998) A tutorial on support vector machines for pattern recognition. Data mining and knowledge discovery 2:121–167

Chaube NR, Lele N, Misra A, et al. (2019) Mangrove species discrimination and health assessment using AVIRIS-NG hyperspectral data. Curr. Sci 116:1136

Chen B, Xiao X, Ye H, et al. (2018) Mapping forest and their spatial–temporal changes from 2007 to 2015 in tropical hainan island by integrating ALOS/ALOS-2 L-Band SAR and landsat optical images. IEEE Journal of Selected Topics in Applied Earth Observations and Remote Sensing 11:852–867

Congalton RG (1991) A review of assessing the accuracy of classifications of remotely sensed data. Remote sensing of environment 37:35–46

Daehler CC (2005) Upper-montane plant invasions in the Hawaiian Islands: patterns and opportunities. Perspectives in Plant Ecology, Evolution and Systematics 7:203–216

de Abreu RC, Durigan G (2011) Changes in the plant community of a Brazilian grassland savannah after 22 years of invasion by Pinus elliottii Engelm. Plant Ecology & Diversity 4:269–278

Delalay M, Tiwari V, Ziegler AD, et al. (2019) Land-use and land-cover classification using Sentinel-2 data and machine-learning algorithms: operational method and its implementation for a mountainous area of Nepal. Journal of Applied Remote Sensing 13:014530

Donnelly J (1985) Community structure of epigaeic ants in a pine plantation and in newly burnt fynbos. Journal of the Entomological Society of southern Africa 48:259–265

Dye P, Jarmain C (2004) Water use by black wattle (Acacia mearnsii): implications for the link between removal of invading trees and catchment streamflow response: working for water. South African Journal of Science 100:40–44

Erinjery JJ, Singh M, Kent R (2018) Mapping and assessment of vegetation types in the tropical rainforests of the Western Ghats using multispectral Sentinel-2 and SAR Sentinel-1 satellite imagery. Remote Sensing of Environment 216:345–354

Foody G, Hill R (1996) Classification of tropical forest classes from Landsat TM data. International journal of remote sensing 17:2353–2367

Gaertner M, Novoa A, Fried J, et al. (2017) Managing invasive species in cities: a decision support framework applied to Cape Town. Biological Invasions 19:3707–3723

George R, Padalia H, Kushwaha S (2014) Forest tree species discrimination in western Himalaya using EO-1 Hyperion. International Journal of Applied Earth Observation and Geoinformation 28:140–149

Gorelick N, Hancher M, Dixon M, et al. (2017) Google Earth Engine: Planetary-scale geospatial analysis for everyone. Remote sensing of Environment 202:18–27

Hu S, Qin J, Ren J, et al. (2020) Automatic Extraction of Water Inundation Areas Using Sentinel-1 Data for Large Plain Areas. Remote Sensing 12:243

Iannelli GC, Gamba P (2019) Urban Extent Extraction Combining Sentinel Data in the Optical and Microwave Range. IEEE Journal of Selected Topics in Applied Earth Observations and Remote Sensing 12:2209–2216

Joshi AA, Sankaran M, Ratnam J (2018) ‘Foresting’the grassland: Historical management legacies in forest-grassland mosaics in southern India, and lessons for the conservation of tropical grassy biomes. Biological Conservation 224:144–152

Kattenborn T, Lopatin J, Förster M, et al. (2019) UAV data as alternative to field sampling to map woody invasive species based on combined Sentinel-1 and Sentinel-2 data. Remote sensing of environment 227:61–73

Laurin GV, Liesenberg V, Chen Q, et al. (2013) Optical and SAR sensor synergies for forest and land cover mapping in a tropical site in West Africa. International Journal of Applied Earth Observation and Geoinformation 21:7–16

Lawrence RL, Wright A (2001) Rule-based classification systems using classification and regression tree (CART) analysis. Photogrammetric engineering and remote sensing 67:1137–1142

Le Maitre DC, Gaertner M, Marchante E, et al. (2011) Impacts of invasive Australian acacias: implications for management and restoration. Diversity and Distributions 17:1015–1029

Ledgard N (2001) The spread of lodgepole pine (Pinus contorta, Dougl.) in New Zealand. Forest Ecology and Management 141:43–57

Lele A, Arasumani M, Vishnudas C, et al. (2020) Elevation and landscape change drive the distribution of a montane, endemic grassland bird. Ecology and evolution 10:7755–7767

Lim J, Kim K-M, Jin R (2019) Tree species classification using hyperion and sentinel-2 data with machine learning in south Korea and China. ISPRS International Journal of Geo-Information 8:150

Lu L, Tao Y, Di L (2018) Object-based plastic-mulched landcover extraction using integrated Sentinel-1 and Sentinel-2 data. Remote Sensing 10:1820

Marshall M, Thenkabail P (2015) Advantage of hyperspectral EO-1 Hyperion over multispectral IKONOS, GeoEye-1, WorldView-2, Landsat ETM+, and MODIS vegetation indices in crop biomass estimation. ISPRS Journal of Photogrammetry and Remote Sensing 108:205–218

McConnachie MM, Wilgen BW, Richardson DM, et al. (2015) Estimating the effect of plantations on pine invasions in protected areas: a case study from South Africa. Journal of Applied Ecology 52:110–118

Melgani F, Bruzzone L (2004) Classification of hyperspectral remote sensing images with support vector machines. IEEE Transactions on geoscience and remote sensing 42:1778–1790

Mitchell AL, Tapley I, Milne AK, et al. (2014) C-and L-band SAR interoperability: Filling the gaps in continuous forest cover mapping in Tasmania. Remote sensing of environment 155:58–68

Mooney HA (2005) Invasive alien species: a new synthesis. Island press

Mountrakis G, Im J, Ogole C (2011) Support vector machines in remote sensing: A review. ISPRS Journal of Photogrammetry and Remote Sensing 66:247–259

Pellikka PK, Lötjönen M, Siljander M, et al. (2009) Airborne remote sensing of spatiotemporal change (1955–2004) in indigenous and exotic forest cover in the Taita Hills, Kenya. International Journal of Applied Earth Observation and Geoinformation 11:221–232

Pu R, Bell S, Meyer C, et al. (2012) Mapping and assessing seagrass along the western coast of Florida using Landsat TM and EO-1 ALI/Hyperion imagery. Estuarine, Coastal and Shelf Science 115:234–245

Raczko E, Zagajewski B (2017) Comparison of support vector machine, random forest and neural network classifiers for tree species classification on airborne hyperspectral APEX images. European Journal of Remote Sensing 50:144–154

Richardson D, Van Wilgen B (1986) Effects of thirty-five years of afforestation with Pinus radiata on the composition of mesic mountain fynbos near Stellenbosch. South African Journal of Botany 52:309–315

Richardson DM (1998) Forestry trees as invasive aliens. Conservation biology 12:18–26

Rodriguez-Galiano VF, Ghimire B, Rogan J, et al. (2012) An assessment of the effectiveness of a random forest classifier for land-cover classification. ISPRS Journal of Photogrammetry and Remote Sensing 67:93–104

Rundel PW, Dickie IA, Richardson DM (2014) Tree invasions into treeless areas: mechanisms and ecosystem processes. Biological Invasions 16:663–675

Salas EAL, Subburayalu SK, Slater B, et al. (2020) Mapping crop types in fragmented arable landscapes using AVIRIS-NG imagery and limited field data. International Journal of Image and Data Fusion 11:33–56

Shaharum NSN, Shafri HZM, Ghani WAWAK, et al. (2020) Oil palm mapping over Peninsular Malaysia using Google Earth Engine and machine learning algorithms. Remote Sensing Applications: Society and Environment:100287

Shimizu K, Ota T, Mizoue N (2019) Detecting Forest Changes Using Dense Landsat 8 and Sentinel-1 Time Series Data in Tropical Seasonal Forests. Remote Sensing 11:1899

Six LJ, Bakker JD, Bilby RE (2013) Loblolly pine germination and establishment in plantations and grasslands of northern Uruguay. Forest ecology and management 302:1–6

Skowronek S, Ewald M, Isermann M, et al. (2017) Mapping an invasive bryophyte species using hyperspectral remote sensing data. Biological Invasions 19:239–254

Slagter B, Tsendbazar N-E, Vollrath A, et al. (2020) Mapping wetland characteristics using temporally dense Sentinel-1 and Sentinel-2 data: A case study in the St. Lucia wetlands, South Africa. International Journal of Applied Earth Observation and Geoinformation 86:102009

Thenkabail PS, Enclona EA, Ashton MS, et al. (2004) Hyperion, IKONOS, ALI, and ETM+ sensors in the study of African rainforests. Remote Sensing of Environment 90:23–43

Veloso A, Mermoz S, Bouvet A, et al. (2017) Understanding the temporal behavior of crops using Sentinel-1 and Sentinel-2-like data for agricultural applications. Remote sensing of environment 199:415–426

Wheeler J, Rodriguez-Veiga P, Balzter H, et al. (2017) Forest mapping of the congo basin using synthetic aperture radar (SAR). Earth Observation for Land and Emergency Monitoring 57

Zalba SM, Cuevas YA, Boó RM (2008) Invasion of Pinus halepensis Mill. following a wildfire in an Argentine grassland nature reserve. Journal of Environmental Management 88:539–546

Zhang W, Brandt M, Wang Q, et al. (2019) From woody cover to woody canopies: How Sentinel-1 and Sentinel-2 data advance the mapping of woody plants in savannas. Remote Sensing of Environment 234:111465

